# Extreme acid resistance of human cheek epithelial cells

**DOI:** 10.1101/2020.01.23.903831

**Authors:** Muhammad Ahmad, Irsa Mateen, Aisha Tahir, Rana Salman Anjum

**Affiliations:** School of Life Sciences, Forman Christian College (A Chartered University), 54600, Lahore, Pakistan; Institute of Biochemistry and Biotechnology, University of the Punjab, 54590, Lahore, Pakistan; Department of Biochemistry, University of Health Sciences, 54000, Lahore, Pakistan

**Keywords:** Cheek epithelial cells, squamous epithelial cells, buccal mucosa, oral cavity, acid exposure, acid resistance

## Abstract

Human cheek epithelial cells, despite residing in an almost neutral environment of the mouth, are exposed to many acidic stimuli, so these cells may possess acid resistance. Our research’s objective is to find out if isolated human cheek epithelial cells can survive in highly acidic conditions, and whether there is any difference in acid resistance between cheek epithelial cells of males and females; and to establish a protocol. We collected 3 buccal swab samples from each male or female participant and gave following 3 different treatments to their samples: (i) Milli-Q water (ii) HCl with pH of −0.48 (iii) HCl with pH of - 0.48 and, then, rehydration with Milli-Q water. Staining and microscopic analysis showed that cheek epithelial cells did not burst after acid treatment. However, they slightly shrunk in size and were stained substantially darker. Moreover, cells that were rehydrated after acid treatment, provided further evidence of their survival because these cells, as a result of rehydration, had substantially regained bigger size as well as lighter staining. Furthermore, at pH of −0.48, no difference was observed in acid resistance of the cells from the two genders. Results of this study, which is the only one to incubate cheek epithelial cells at pH < 2.0, manifest that the human cheek epithelial cells, isolated from any of the two genders, are extremely acid-resistant and do not burst in highly acidic conditions. Besides, these cells may also be useful as model cells for the study of acid-resistant pathogens and acid-resistant cancer cells.

## Introduction

Human cheek epithelial cells are present in the inner lining of the cheek, i.e. part of buccal mucosa (Shum et al. 2018). The epithelium layer of the buccal mucosa is stratified squamous epithelium in nature (Slaughter 1953), and cheek epithelial cells, usually observed in a light microscope, are squamous epithelial cells (Dawidson 2015). Human cheek epithelial cells are bathed with saliva that has a pH of 6.6-7.1 (Zhang et al. 2016). Moreover, the mean pH value for the soft and hard palate, different parts of the tongue, buccal mucosa, and floor of the mouth is also close to neutral, i.e. ~6.78 (Aframian et al. 2006). Thus, it may be inferred that cheek epithelial cells can survive only at neutral pH, i.e. pH of their usual environment, and will burst in response to very low pH values. However, most of our natural food is acidic in nature, and foods like yogurt, vinegar, pickled vegetables, and most fruits are highly acidic (pH < 4.6) in nature (McGlynn 2003). Moreover, lemon juice can have a pH of 2.0 (Healthline n.d.), and Coca Cola has a pH of 2.37 (Reddy et al. 2016). Furthermore, gastric contents, having a pH of nearly 1, can reach an oral cavity during gastroesophageal reflux (though that usually does not cause much reduction in pH of the oral cavity which drops to just ~6.65) (Herregods et al. 2015). In addition, the oral cavity of a person can be exposed to acids of varying concentrations and pH values, which can be very extreme in some cases, when he/she ingests acid accidentally or for suicide (Marsella et al. 2017). Other acidic stimuli, which can affect the oral cavity, also include methamphetamine drugs that can have pH as low as 3.02 (Grobler et al. 2011). These acidic stimuli cause erosion of enamel and dentine, and our cheek epithelial cells are also exposed to them. Sometimes, these acidic stimuli can exceed the buffering capacity of saliva, especially in people suffering from hyposalivation that do not have much saliva for buffering (Grobler et al. 2011; Ranjitkar et al. 2012). Thus, if cheek epithelial cells are not resistant to low pH conditions, then the acidic stimuli mentioned above, which are frequently encountered by us, can cause damage to our cheek epithelial cells.

However, the epithelial cells from esophageal and laryngeal mucosa, despite living in a neutral pH environment (Campagnolo et al. 2014; Tutuian and Castell 2006), have been shown to be acid-resistant (but not resistant to pepsin and bile) in the following studies. Salo et al. (1983) tested the effect of acidic conditions on the esophagus of rabbits. They isolated 6-7 cm of the esophageal tube from the digestive tract of the rabbits, and treated it with 150 mM HCl. Even after 3.5 hours of incubation, no prominent changes occurred in the mucosa and submucosa of the esophagus. Orlando (2010) stated that, during the Bernstein test, the esophagus of healthy people could tolerate 30 minutes of perfusion with HCl of pH 1, and it did not cause heartburn in those people. Hopwood et al. (1981) also studied the effect of 0.1 N HCl on biopsies of the human esophageal mucosa. They found that there was very little damage caused, by 0.1 N HCl, to the esophageal mucosa. In addition, the cells of esophageal mucosa remained unchanged and intact. Nonetheless, gastric juice at pH of 1-3 severely affected the cells of the esophageal mucosa. Herregods et al. (2015) stated that the acid alone causes less damage during gastroesophageal reflux, and major damage is caused by pepsin and bile components. Besides, according to many studies including those done by Calabrese et al. (2003) and Villanacci et al. (2001), gastric reflux is widely known to cause dilation of intercellular spaces of esophageal tissue, and this dilation of intercellular spaces of esophageal tissue is regarded as a very sensitive marker of gastroesophageal reflux (Calabrese et al. 2003; Villanacci et al. 2001). In addition, Franchi et al. (2007) reported that similar to acid reflux-caused dilation of intercellular spacing observed in esophageal mucosa, acid reflux also caused dilation of intercellular spacing in laryngeal mucosa of patients suffering from laryngopharyngeal reflux (which is considered another name for acid reflux or gastroesophageal reflux). However, no gross change was observed in the laryngeal mucosa of these patients’ tissues and their squamous epithelial cells remained intact after they have suffered acid reflux having a pH of 1-2. Furthermore, Campagnolo et al. (2014) treated different parts of porcine laryngeal mucosa with buffers of pH 4.0 and 2.0. They found that those parts of laryngeal mucosa that contained columnar epithelium were sensitive to both buffers (having pH values of 4.0 and 2.0), while the parts containing squamous epithelium were resistant to both buffers (having pH values of 4.0 and 2.0). Hence, squamous epithelial cells from both esophageal and laryngeal mucosae are resistant to very low values of pH.

Although no study was found in the existing literature that focused on acid resistance of individual cheek epithelial cells or buccal mucosa, some studies were found that, for analysis of transport of different substances across buccal mucosa/epithelium, incubated buccal mucosa/epithelium in low pH conditions that indirectly pointed to acid resistance of buccal mucosa. Birudaraj et al. (2005) incubated porcine buccal mucosa at acidic pH values with the lowest pH of 3.0 while checking the effect of pH on the transport of buspirone across the buccal mucosa. Gandhi and Robinson (1991) and Dowty et al. (1992) used rabbit buccal mucosa in diffusion cells and, for this purpose, they incubated rabbit buccal mucosa at different acidic pH values with the lowest pH values of 4.0 and 2.0, respectively.

Thus, many studies described the acid resistance of epithelial cells from esophageal and laryngeal mucosae, but no study focusing on acid resistance of cheek epithelial cells was found. Moreover, no study was found to treat cheek epithelial cells at pH < 2.0 and squamous epithelial cells from any of buccal, esophageal, or laryngeal mucosae, from any organism, at pH < 1.0. Furthermore, calf erythrocytes and human erythrocyte ghosts have been shown to burst in < 5 minutes when incubated at pH of 3.2 (Ivanov 1999). Hence, in this study, we aim to see if isolated cheek epithelial cells can remain intact at very low pH of –0.48 for 30 minutes, and whether there is any difference in the response of cheek epithelial cells of males and females to these extremely acidic conditions.

## Materials and methods

### Participants

10 participants, consisting of 6 male and 4 female participants, were included in this study. All participants belonged to Forman Christian College (A Chartered University). 3 buccal swab samples were taken from each participant with informed consent.

### Smear formation and treatment with HCl

Clean cotton buds were rubbed gently on the inner linings of both left and right cheeks. Microscopic slides were marked (on the underneath surface) to allow clear recognition of the area (throughout the experiment) on which buccal smears would be formed. 3 smears, of inner linings of cheek cells, were formed (each on pre-marked 1cm^2^) on microscopic slides using these cotton buds such that 2 smears were formed on one microscopic slide while the 3^rd^ one was formed on another microscopic slide.

#### 1^st^ microscopic slide

##### i. Smear 1 (buccal swab smear treated with Milli-Q water)

20 μl of Milli-Q water was placed on the smear using a micropipette and spread on the smear using a micropipette tip. After 30 minutes of incubation time, the slide was tilted at an angle of 45°, so that epithelial cells remained adsorbed to the slide while most of the water flowed to the lower end of smear, and this water was absorbed using a paper towel.

##### ii. Smear 2 (buccal swab smear treated with 9.25% HCl)

20 μl of 9.25% HCl was placed on the smear using a micropipette and spread on the smear using a micropipette tip. After an incubation time of 30 minutes, the slide was tilted, and HCl was absorbed as described above in the case of smear 1.

#### 2nd microscopic slide

##### i. Smear 3 (buccal swab smear treated with 9.25% HCl followed by treatment with Milli-Q water)

Initially, a procedure identical to that of smear 2 was given (20 μl of 9.25% HCl was dropped on the smear using a micropipette and spread on the smear using a micropipette tip. 30 minutes of incubation time was given and, then, the slide was tilted and HCl was absorbed by a paper towel as described above in the case of smear 1).

It was followed by rehydration of smear 3 with Milli-Q water. Rehydration was performed by dropping 20 μl of Milli-Q water on acid-treated smear using a micropipette, spreading Milli-Q water on smear using a micropipette tip, and giving an incubation time of 2 minutes. Slide was tilted and liquid was removed by paper towel as described above in case of smear 1. Rehydration was performed 2 more times on smear 3.

### Staining and microscopic observation of slides

Staining was done by adding 10 μl of methylene blue solution, using a micropipette, on 3 smears of each participant (obtained as a result of the above treatment steps). A separate micropipette tip was used to spread a methylene blue solution over each smear, and an incubation time of 3 minutes was given. Then, the slide was tilted at an angle of 45°, so that epithelial cells remained adsorbed to the slide while most of the dye solution flowed to the lower end of smear, and this dye solution was absorbed using paper towels. Microscopic slides were left to air dry for 2 minutes before cover slips were placed on top of smears. These smears were, then, observed by a microscope under 10X eyepiece combined with 100X objective lens (using immersion oil), and photographs of epithelial cells, near the center of smears, were taken by using camera.

## Results

### Microscopic analysis of human cheek epithelial cells

All smears from all participants retained a sufficient number of cheek epithelial cells, after different treatments, for microscopic analysis. After 30 minutes of incubation in 9.25% HCl (that had pH of −0.48), these cells were able to survive, but they underwent slight shrinkage and were stained darker (in comparison to the cells incubated in Milli-Q water for 30 minutes). The cells that were rehydrated, after 30 minutes of incubation with 9.25% HCl, had larger sizes and were stained lighter (similar to the cells incubated with just Milli-Q water for 30 minutes). Photomicrographs of cells treated with all 3 different treatments, from all participants, are presented in Figure 1. The smears have been taken from healthy individuals with no reports of mouth disease. The microscopy and staining was performed following a study published in Scientific Reports by Theda et al., 2018. They have also used a low cost method to stain the cheek cells and visualization was done on light microscope. However, we have used the same method with slight modifications. Green colour arrows indicate the nucleus inside the cell which is stained normal in both A and C panels. While in the B panel it is difficult to visualize the nucleus due to darkly stained cytoplasm. Cheek cells are very delicate cells which get burst and release out the nucleus upon physical shearing. But we made sure throughout the process that no physical shearing happens. Moreover, acids can also damage the cell membranes but it is tempting to speculate that human cheek epithelial cells have mechanism responsible for fighting high concentration of protons. These cells not only uptake protons but also release water through osmotic imbalance that would possibly be a strategy to decrease the acidity inside buccal cavity.

**Figure 1:**
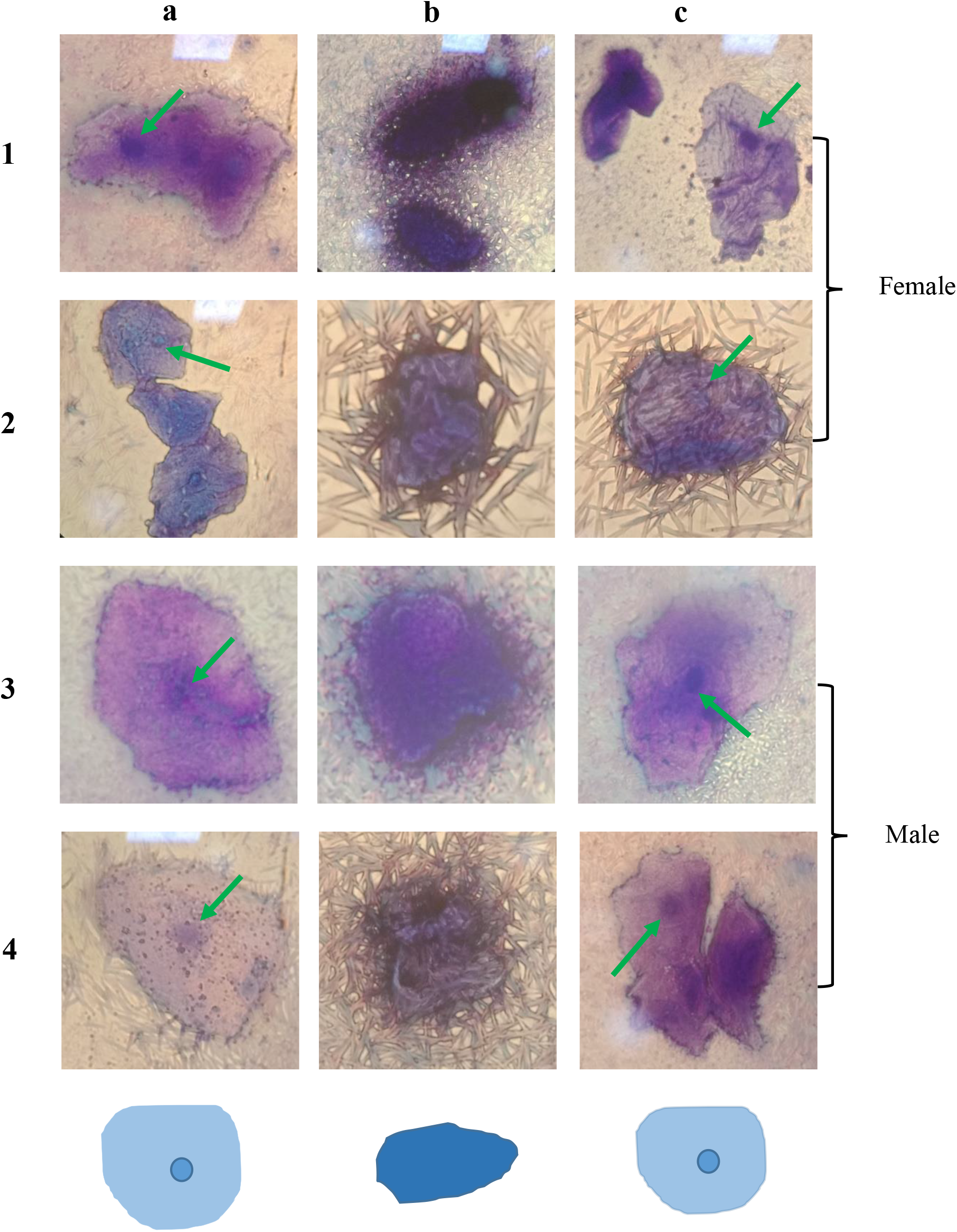
Photomicrographs of isolated human cheek epithelial cells, from individuals, exposed to 3 different treatments. Individuals 1 & 2 were females, while individuals 3 & 4 were males. Treatment A: Milli-Q water for 30 minutes; Treatment B: HCl (with a pH of –0.48) for 30 minutes; Treatment C: Treatment B followed by 3 rounds of rehydration with Milli-Q water. Green arrows indicate a healthy nucleus. Microscopic reconstitution of the cells is placed at the bottom.

## Possible mechanism of pH homeostasis by cheek epithelial cells

In Figure 2, the normal cheek epithelial cell is in equilibrium with water. While the equilibrium is disturbed in the presence of a strong acid, allowing more H^+^ ions to go inside while water molecules are released out of the cell resulting in shrinking. The equilibrium is reversed again in the presence of water and the cell regains its normal shape. This mechanism illustrates the possible role of human cheek epithelial cells in regulating homeostasis of ions as well as defense against an acid stress. The darker staining is possibly due to accumulation of lipid and protein content hence retaining more stain. The nucleus is not visible in darkly stained cell because of the same level staining of cytoplasm surrounding the nucleus. Moreover, higher concentration of H^+^ ions inside the cell would affect the protein metabolism strongly due to charge effect on proteins which may result in cell division arrest. Studies on HeLa cells have shown that acid effects the cell growth but they have tested the effect of pH on HeLa cells close to physiological pH conditions.

**Figure 2:**
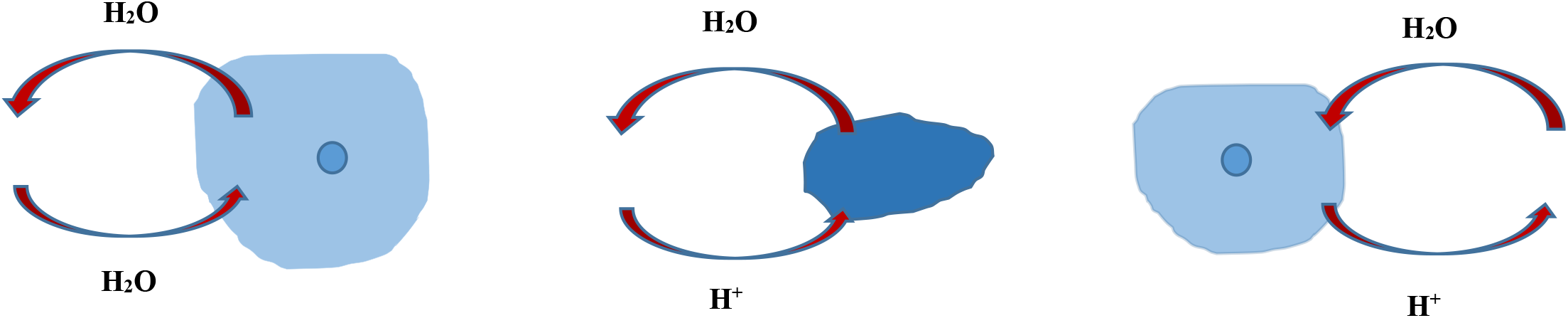
Model of pH homeostasis by cheek epithelial cells. The schematic representation of microscopic reconstitution shows the behavior of human cheek cells in different environmental conditions.

## Discussion

We performed this study to know if cheek epithelial cells can survive in highly acidic conditions, i.e., 30 minutes in 9.25% HCl having a pH of −0.48. Cells incubated for 30 minutes in Milli-Q water were taken as control. These were compared with cells incubated in 9.25% HCl for 30 minutes and with cells rehydrated after 30 minutes of incubation with 9.25% HCl. As compared to the control group, 30 minutes of incubation in acid caused cells to slightly shrink and get stained darker. However, the cheek epithelial cells were able to survive at this extremely low pH (indicated by an intact plasma membrane and a visible nucleus). To confirm the survival of these cells at this pH, after 30 minutes of incubation in 9.25% HCl, we performed 3 rounds of rehydration on cells (by the procedure described in “Methods”). This rehydration treatment allowed the cells to regain larger size and lighter staining that were comparable to the control group. These cells also had intact membranes and a visible nucleus. Thus, it confirmed that human cheek epithelial cells were capable of surviving, for 30 minutes, at a pH of −0.48.

In this study, we devised a new procedure, which was very fast and easy to perform, for testing the effect of the acidic solution on cheek epithelial cells. This procedure took advantage of the observation that if microscopic slide was tilted, then cheek epithelial cells, suspended in HCl (or water), adsorbed on the surface of microscopic slide while HCl (or water) flowed down towards the bottom end of slide, and this HCl (or water) can be absorbed by using an absorbent material like paper towel. Although this procedure looks overly simple, it can be applied for fast testing of effects of acid on any cell type (such as acid-resistant cancer cells and acid-resistant pathogens) capable of adsorbing to microscopic slides. Thus, this procedure can be done on just a microscopic slide, without using a separate container for incubating cells with acid solution, and, therefore, will be easy and time saving in case of a large number of cell/tissue samples.

Moreover, results of our study have shown that, similar to the acid resistance of squamous epithelial cells of esophageal mucosa (Hopwood et al. 1981; Orlando 2010; Salo et al. 1983) and laryngeal mucosa (Campagnolo et al. 2014; Franchi et al. 2007), squamous epithelial cells from buccal mucosa (called cheek epithelial cells) also exhibit acid resistance. Additionally, we have shown resistance of squamous epithelial cells, from buccal mucosa, at a pH value (of −0.48) that is far lower than pH values used in studies on esophageal mucosa (Hopwood et al. 1981; Orlando 2010; Salo et al. 1983) and laryngeal mucosa (Campagnolo et al. 2014; Franchi et al. 2007). However, we have just performed light microscopy analysis (at 1000x), and electron microscopy analysis is expected to reveal more about the acid resistance of cheek epithelial cells. More advanced light microscopy, or electron microscopy, of the three sets of cells that we used in our experiment (normal cells, acid-treated cells, and cells rehydrated after acid treatment) will tell whether cheek epithelial cells are resistant to low pH values at the subcellular levels, e.g., whether membranes of individual organelles burst due to acid treatment (which cannot be observed by a simple light microscope). Importantly, cell culturing will be the best method to test whether these acid-treated cells are totally functional and alive.

Additionally, according to results from our study, cheek epithelial cells should not be damaged by the acidity of the acidic stimuli mentioned above, i.e., gastric contents (during gastroesophageal reflux), acidic foods, acidic drinks, and methamphetamine drugs. It is because the pH values of all of these acidic stimuli are very higher (Grobler et al. 2011; Healthline n.d.; McGlynn 2003; Ranjitkar et al. 2012; Reddy et al. 2016) than the pH of −0.48 that cheek epithelial cells are able to tolerate in our study. Accidental acid intake, causing the pH of buccal mucosa to drop to up to −0.48, is not expected to harm/lyse cheek epithelial cells. Even in case of hyposalivation patients, which do not have much saliva for buffering acid-intakes (Grobler et al. 2011; Ranjitkar et al. 2012), cheek epithelial cells are expected to stay intact upon exposure to the acidic stimuli mentioned above because, in our study, isolated cheek epithelial cells were suspended in acid, so no or trace amounts of saliva were present for buffering action. Besides, our study also supports the stand of (Herregods et al. 2015) that, during gastroesophageal reflux, acid alone does not cause damage to squamous epithelial cells, and other components of gastric juice are responsible for most of the damage. Moreover, it has been shown that the cheek epithelial cells collapse because of osmolarity differences (Lee et al. 1994), however, we present here our first ever report that the cheek cells are resistant to strong acidic environment. These cells shrink but do not collapse. Indeed, further research will be required to investigate the mechanism of acid stability.

## Conclusion

Our study has shown that isolated human cheek epithelial cells are resistant to pH of −0.48, for 30 minutes, so ingestion of acid having pH equal to or higher than −0.48 is not expected to directly damage/lyse cheek epithelial cells. We do not know the effect of ingestion of acid having pH lower than −0.48, on individual cheek epithelial cells because we have not checked the effect of pH values lower than −0.48 on isolated cheek epithelial. Therefore, testing the effect of even lower pH values (than −0.48) on cheek epithelial cells, which will require more carefully controlled conditions, will provide information about the limits of acid resistance of human cheek epithelial cells, i.e., the lowest pH value that can be tolerated by human cheek epithelial cells. Moreover, in the future, acid-treated human cheek epithelial cells should be cultured to see if they can reproduce or not. Furthermore, checking the effect of basic pH or excessive heat on human cheek epithelial cells will provide more information about the resistance of these cells. Knowledge and guidelines that are obtained from this research can pave the way for anybody interested in a more detailed exploration of acid resistance of human cheek epithelial cells and, due to their easy availability, these human cheek epithelial cells can also be adopted as model cells for studying and countering acid resistance of acid-resistant pathogens and acid-resistant cancer cells.

## Acknowledgements

We would like to thank School of Life Sciences, Forman Christian College (A Chartered University) for providing support to conduct this study. We are also thankful to all volunteers for providing samples.

## Declaration of informed consent

The authors declare that samples were taken from each participant with informed consent and that the study was approved from Ethical Review Committee of Forman Christian College.

